# Characterization of astrocyte density in the Pitt-Hopkins Syndrome mouse model of ASD

**DOI:** 10.1101/2025.07.10.664109

**Authors:** Sara M. Stump, Joseph F. Bohlen, BaDoi Phan, Brady J. Maher

## Abstract

Transcription factor 4 (TCF4) is a proneural basic helix-loop-helix transcription factor that plays a critical role in brain development and is associated with a variety of psychiatric disorders including autism spectrum disorder (ASD), major depressive disorder, and schizophrenia. Autosomal dominant mutations in *TCF4* result in a profound neurodevelopmental disorder called Pitt-Hopkins Syndrome (PTHS). Germline TCF4 loss-of-function (LOF) studies using human and mouse models have identified dysregulation in neural cell proliferation, genesis, and specification which lead to disruption in neuronal, astroglial and oligodendroglial lineages. In this study, we focused on the role of TCF4 in the genesis of the astrocyte lineage, specifically in the context of modeling PTHS. We show that germline heterozygous mutations in *Tcf4* had no effect on the expression of astrocyte marker genes in primary astrocyte cultures and whole brain lysates. Immunohistochemical (IHC) analysis of pan- and subclass-specific astrocyte markers showed *Tcf4* mutation had no effect on the proportions of astrocytes in the dorsal cortex and corpus callosum. Lastly, we tracked ventrally-derived astrocytes using an Nkx2.1 reporter mouse and observed that germline *Tcf4* LOF did not result in misallocation of ventrally-derived astrocytes into the dorsal cortex, a phenotype previously observed when both *Tcf4* alleles were conditionally deleted in the Nkx2.1 lineage. These data indicate that germline heterozygous TCF4 LOF, which models PTHS, does not appear to affect the astrocyte lineage at the cell population level.

## INTRODUCTION

ASD is a highly heritable, common childhood disorder, reported to occur between 3.4 to 62.6 out of every 1,000 children^1,2^ Core features of ASD include impaired social interaction, delayed or absence of language, stereotypical patterns of behavior, and can present with other comorbidities such as intellectual disability, epilepsy, anxiety, depression, attention deficits, and sleep disorders^3^. The majority of ASD cases are thought to stem from common variant risk^4,5^, while approximately 10% of cases are driven by rare and *de novo* variants that confer substantial risk^6,7^. One syndromic form of ASD is Pitt-Hopkins syndrome (PTHS), which is caused by loss-of-function mutations (LOF) in the *TCF4* gene (not *TCF7L2*, which encodes T-Cell Factor 4). PTHS is a rare neurodevelopmental disorder characterized by intellectual disability, failure to acquire language, deficits in motor learning, hyperventilation, gastrointestinal abnormalities and autistic behavior ^8^. TCF4 is highly expressed in dorsal and ventral neuroprogenitor cells in the developing CNS, where it is shown to regulate proliferation, differentiation, and specification ^9–11^. Heterozygous *Tcf4* LOF, which models PTHS, was shown to alter the proportions of upper-layer excitatory neurons, distinct classes of inhibitory neurons, and the oligodendroglial lineage ^9,12–15^. *Tcf4* also appears to regulate astroglia, as germline homozygous *Tcf4* LOF resulted in a severe depletion of midline glia (GFAP+) in the indusium griseum and callosal wedge ^10^, as well as conditional homozygous deletion in the Nkx2.1 lineage resulted in ectopic allocation of ventrally derived astrocytes in the dorsal neocortex ^16^.

Astrocytes are a heterogeneous subset of glial cells that play diverse functional roles in the central nervous system, and their dysregulation is associated with several neurological and neuropsychiatric disorders. Representing 20-40% of cells within the human neocortex, astrocytes perform a multitude of functions. They participate in synaptic function by maintaining ion and neurotransmitter homeostasis, providing metabolic support, and releasing gliotransmitters which can modulate synaptic transmission ^17^. Their direct communication with neurons leads to an essential role for astrocytes in brain development, contributing to synaptogenesis, synaptic development, axon guidance, and synapse and axon pruning ^18–20^.

In this study, we focused on the role of *Tcf4* in the regulation of astrocyte development in the context of PTHS. Specifically, we asked if germline heterozygous *Tcf4* mutation (*Tcf4^+/tr^*) alters astrocyte marker gene expression and/or the density of astrocytes in the dorsal cortex. We show in primary astrocyte cultures and whole brain samples that *Tcf4* mutation had no appreciable effect on astrocyte marker gene expression. Similarly, *Tcf4* mutation had no effect on the total density of astrocyte populations in the late adolescent cortex or the density of fibrous (GFAP+) or protoplasmic (s100β) subclasses of astrocytes. Lastly, we show that germline *Tcf4* mutation had no effect on the density of ventrally-derived astrocytes populations residing in the dorsal cortex. Together, these results suggest germline *Tcf4* mutation mimicking PTHS does not appear to alter astrocytes at the population level in the murine dorsal cortex.

## MATERIALS AND METHODS

### Animals and tissue collection

The *Tcf4^+/tr^* mouse model of PTHS is heterozygous for an allele encoding deletion of the DNA-binding domain of *TCF4* (B6;129-TCF4tm1Zhu/J, stock #013598, Jackson Laboratory). TdTom listed as B6.Cg-Gt(ROSA) 26Sortm14(CAG-tdTomato)Hze/J (Jackson Laboratory stock #007909). Nkx2.1 Cre listed as C57BL/6J-Tg(Nkx2-1-cre)2Sand/J (Jackson Laboratory stock #008661). Mouse colonies were backcrossed for at least six generations in the C57/B6 background, maintained by The Lieber Institute for Brain Development’s Animal Facility on a 12h light/dark cycle and fed ad libitum. *Tcf4^+/tr^*mouse samples were matched with samples from *Tcf4^+/+^* (WT) littermates, and sex was randomly selected in each genotype and age group. All procedures were performed in accordance with the National Institutes of Health Guide for the Care and Use of Laboratory Animals and approved by the Johns Hopkins University School of Medicine’s Institutional Animal Care and Use Committee.

### Immunohistochemistry

Mice (postnatal days 28 and 42) were anesthetized with isoflurane and perfused with cold PBS followed by 4% paraformaldehyde (4% PFA). Brains were dissected out and post-fixed in 4% PFA at 4°C overnight before transferring to PBS. Sections were sliced to a thickness of 55 µm using the Leica VT1000S vibratome. Slices were washed three times with 0.4% Tween-20 and blocked in goat serum (10%) for 2h at room temperature on an orbital shaker. After blocking, primary antibody was added to goat serum (2%) in 0.4% Tween-20 and slices were incubated at 4°C overnight. The following day, slices were washed three times with 0.4% Tween-20 and incubated with secondary antibody for 2h at room temperature. Following incubation, slices were washed three times with 0.4% Tween-20 before counterstain with DAPI (Invitrogen, D1306). Visualization was carried out on a ZEISS LSM 700 Confocal. Two images per animal were averaged for cell counts. Cell counts were compared between genotypes using two-sided unpaired *t*-tests. Below is a list of antibodies used in this study.

#### IHC Antibodies

SOX9 1:250 rabbit HPA001758 Millipore Sigma

ALDH1L1 1:500 mouse WH0010840M1 Millipore Sigma

GFAP 1:500 chicken ab4674 abcam

s100β 1:500 chicken S100B-0100 aveslabs

Alexa Fluoro 488 Goat anti-Chicken 1:1000 Invitrogen A11039

Alexa Fluoro 488 Goat anti-Mouse 1:1000 Invitrogen A11001

Alexa Fluoro 488 Goat anti-Rabbit 1:1000 Invitrogen A11008

Alexa Fluoro 546 Goat anti-Rabbit 1:1000 Invitrogen A11010

Alexa Fluoro 647 Goat anti-Chicken 1:1000 Invitrogen A32933

Alexa Fluoro 633 Goat anti-Rabbit 1:1000 Invitrogen A21071

### Primary astrocyte cultures

Primary astrocytes were prepared as previously described with modifications^21,22^. Briefly, cortices were extracted, and meninges removed from postnatal day 2–3 pups. After dissection cortices were triturated with a 1 ml pipette 10 times to break down the tissue into smaller pieces. The tissue was then incubated with 300μl 2.5% trypsin-EDTA (Gibco) and 100μl DNase (100μg/ml; Sigma) on a shaker at 150 rpm at 37°C for 30–45 minutes. Subsequently, the cells were passed through a 40μm mesh filter, and plated in Glial media (MEM, 5% FBS, Glucose (20mM), Sodium bicarbonate (0.2mg/ml), L-Glutamine (2.5mM) and 1% Pen/Strep). After 24h, the plates were vigorously tapped and shaken to remove any loosely attached cells. The cells were cultured for 6 days with medium changes every other day before collection in TRIzol (Life Technologies) for RNA extraction.

### RNA isolation and qPCR

The transcription expression of genes of interest were measured in whole mouse brains at postnatal day 42 and in DIV6 primary astrocyte cultures. Samples were first collected and homogenized using TRIzol (Life Technologies). Total RNA was extracted using phenol:chloroform isolation followed by purification using the RNeasy Mini Prep Kit (Qiagen 74104). Complementary DNA (cDNA) was prepared from purified RNA using the High-Capacity RNA-to-cDNA Kit (Applied Biosystems, 4387406). qPCR assays were carried out using the QuantStudio3 Real-Time PCR system (Applied Biosystems). qPCR was performed according to the manufacturer’s instructions using Taqman probes. Data are expressed as the fold change of the gene of interest normalized to *Gapdh* expression using 2^-ΔΔCT^. Fold change values were compared between genotypes using two-sided unpaired *t*-tests. Below is a list of Taqman probes used in this study.

#### Taqman probes

SOX9 Thermo Fisher Scientific Mm00448840_m1

GFAP Thermo Fisher Scientific Mm01253033_m1

ALDH1L1 Thermo Fisher Scientific Mm03048957_m1

NDRG2 Thermo Fisher Scientific Mm00443483_m1

FgFr3 Thermo Fisher Scientific Mm00433294_m1

Gja1 Thermo Fisher Scientific Mm01179639_s1

Slc1a3 Thermo Fisher Scientific Mm00600697_m1

Gapdh Thermo Fisher Scientific Mm99999915_g1

### scRNA-seq analysis

Single-cell RNA sequencing data from human dorsolateral prefrontal cortex samples was obtained from the PsychENCODE brainSCOPE dataset from http://brainscope.psychencode.org (PMID 38781369). Biospecimens with fewer than 10 confidently annotated cell types were excluded. Gene expression was pseudobulked by cell type across biological replicates and log-normalized. TCF4 expression distributions were visualized using violin plots, with a reference line indicating mean astrocyte expression.

## RESULTS

To begin to investigate the role of TCF4 in astrocyte development, we first investigated cell-type specific expression of *TCF4* from publicly available single-cell RNA sequencing (scRNA-seq) data from human postmortem dorsolateral prefrontal cortex ^23^. We observed that TCF4 was moderately expressed in astrocytes compared to all other classified cell-types (Figure 1A). We next analyzed expression of astrocyte marker genes in primary astrocyte cultures and whole brain samples obtained from WT and *Tcf4* heterozygous (*Tcf4^+/tr^*) littermates (Figure 1B). We performed qPCR on DIV6 cultures and observed no difference in expression of *Sox9*, *Aldh1l1*, *Gfap*, *Ndrg2*, *Fgfr3*, *Gja1*, and *Slc1a3* between genotypes (Figure 1C). Similarly, we observed no differences in astrocyte marker expression from P42 brain lysates between *Tcf4* genotypes (Figure 1D).

**Figure 1:**
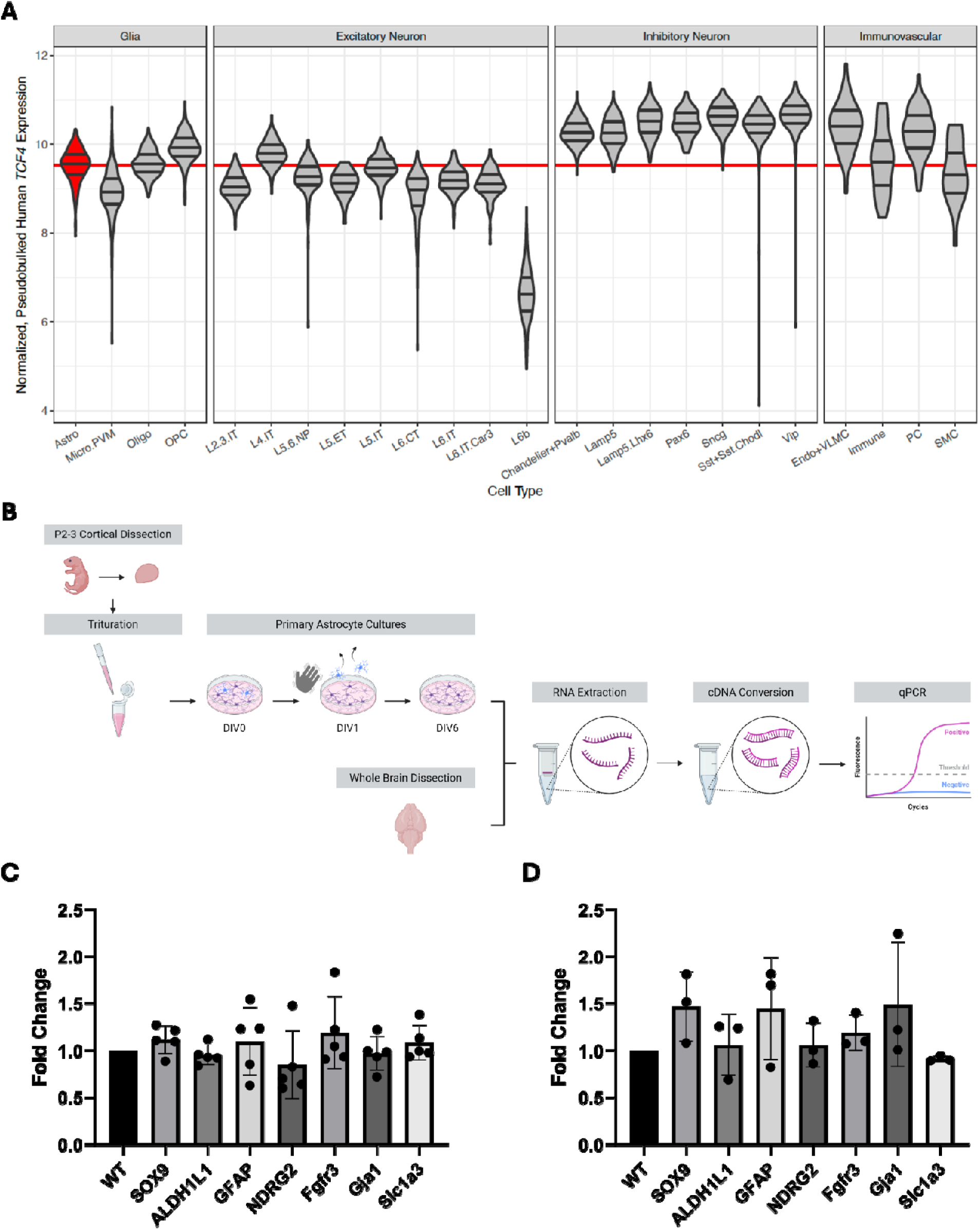
Astrocyte transcript expression is unimpaired in the PTHS mouse model. **(A)** *TCF4* expression across brain cell types in human dorsolateral prefrontal cortex. Violin plots showing normalized, pseudobulked *TCF4* expression across major brain cell types grouped by class (Glia, Excitatory Neuron, Inhibitory Neuron, Vascular). Red line marks the average *TCF4* expression in astrocytes. Violin plots display quartile lines. (**B**) Graphical representation of the astrocyte plating and whole brain dissections for qPCR experiments. (**C**) qPCR of astrocyte-specific gene expression in DIV6 primary astrocyte cultures. Expression of *Sox9* (*P* = 0.3866), *Aldh1l1* (*P* = 0.6696), *Gfap* (*P* = 0.8143), *Ndrg2* (*P* = 0.6133), *Fgfr3* (*P* = 0.5536), *Gja1* (*P* = 0.8409), and *Slc1a3* (*P* = 0.5782) was not different between Tcf4^+/tr^ and WT cultures (*Tcf4^+/tr^ n* = 5 mice; WT *n* = 2 mice). (**D**) qPCR of astrocyte-specific gene expression in P42 whole brain lysates of *Tcf4^+/tr^*and WT littermates. Expression of *Sox9* (*P* = 0.1901), *Aldh1l1* (*P* = 0.9157), *Gfap* (*P* = 0.3716), *Ndrg2* (*P* = 0.7276), *Fgfr3* (*P* = 0.3281), *Gja1* (*P* = 0.2965), and *Slc1a3* (*P* = 0.6308) was not different between genotypes (*Tcf4^+/tr^ n* = 3 mice; WT n = 3 mice).

To quantify the overall population density of astrocytes within the gray and white matter, we performed immunohistochemical (IHC) quantification on coronal brain sections containing the cortex and corpus callosum (CC) from P42 *Tcf4^+/tr^* and WT littermates. To control for brain regional heterogeneity of astrocyte density, all analysis was performed on coronal sections containing the dorsal hippocampus (Figure 2A). Astrocytes were identified by their expression of the pan-astrocyte marker SOX9 and cell counts were normalized with DAPI (Figure 2). The overall astrocyte population (SOX9+/DAPI) was not different between *Tcf4^+/tr^* and WT littermates in both the cortex (Figure 2B,C) and CC (Figure 2D,E).

**Figure 2:**
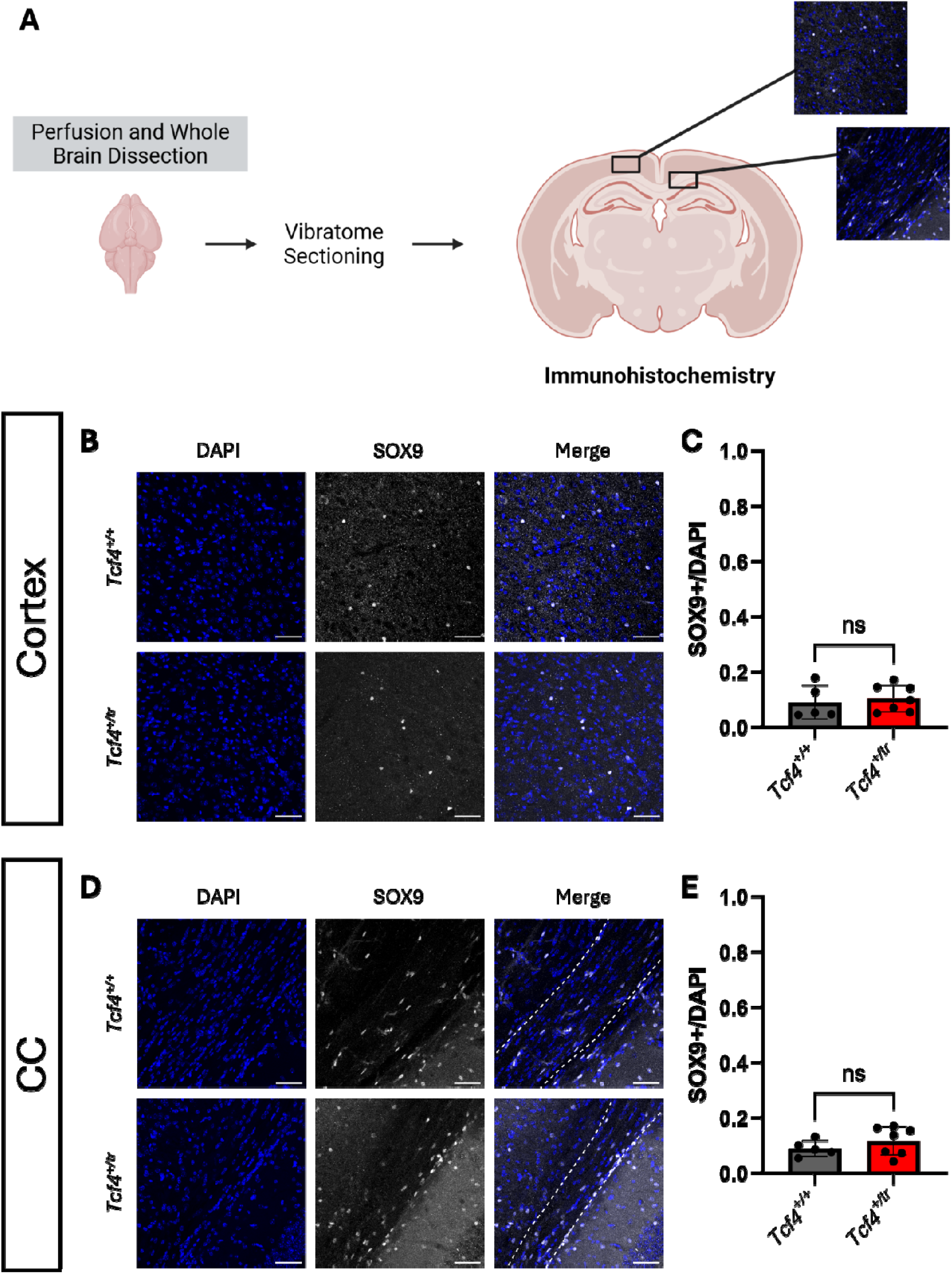
Total astrocyte populations are unimpaired in the PTHS mouse model. (**A**) Graphical representation of the cortex and CC regions quantified using IHC. (**B**) IHC of the overall astrocyte population in the adult cortex of *Tcf4^+/tr^* and WT littermates. Tissue stained for DAPI (blue, column 1) and SOX9 (gray, column 2). (**C**) Summary plot showing no change in the overall proportion of SOX9+DAPI+/DAPI+ cells between *Tcf4^+/tr^* and WT mice in the cortex (*Tcf4^+/tr^* 0.1037 ± 0.01860 *n* = 7 mice, WT 0.08971 ± 0.02735 *n* = 5 mice; *P* = 0.6698). (**D**) IHC of the overall astrocyte population in the adult CC of *Tcf4^+/tr^* and WT littermates. Tissue stained for DAPI (blue, column 1) and SOX9 (gray, column 2). (**E**) Summary plot showing no change in the overall proportion of SOX9+DAPI+/DAPI cells between *Tcf4^+/tr^* and WT mice in the CC (*Tcf4^+/tr^* 0.1162 ± 0.01963 *n* = 7 mice, WT 0.08882 ± 0.01316 *n* = 5 mice, *P* = 0.3155). Data are presented as mean values ± SEM, scale bars 50 µm.

To further investigate protoplasmic and fibrous subpopulations of astrocytes in the *Tcf4^+/tr^* mouse model, we performed IHC for ALDH1L1, GFAP, and s100β. ALDH1L1 identifies a large proportion of the overall astrocyte population, including astrocytes in cortical gray matter, the CC, and in both GFAP+ and GFAP-astrocytes ^24^. We quantified the ALDH1L1+ astrocyte population (ALDH1L1+SOX9+/SOX9+) in the cortex and CC of P42 *Tcf4^+/tr^* and WT littermates and observed no difference between *Tcf4^+/tr^* and WT littermates in both the cortex (Figure 3A, B) and CC (Figure 4A, B). GFAP is a marker for white matter fibrous astrocytes and reactive astrocytes ^25^. The GFAP+ fibrous astrocyte populations (GFAP+SOX9+/SOX9+) were not different by genotype in the cortex (Figure 3C, D) and CC (Figure 4C, D). s100β identifies protoplasmic astrocytes within cortical gray matter and a subset of myelinating OLs ^25,26^. To preclude counting s100β+ OLs, we only counted s100β+SOX9+ double positive cells. Protoplasmic astrocyte populations (s100β+SOX9+/SOX9+) in the gray matter cortex and CC were also no different between *Tcf4^+/tr^* and WT littermates (Figure 3E, F; Figure 4E, F). Together, these results indicate that astrocyte marker expression and the density of the overall astrocyte population, including specific subpopulations, do not appear to be dysregulated in the PTHS mouse model.

**Figure 3:**
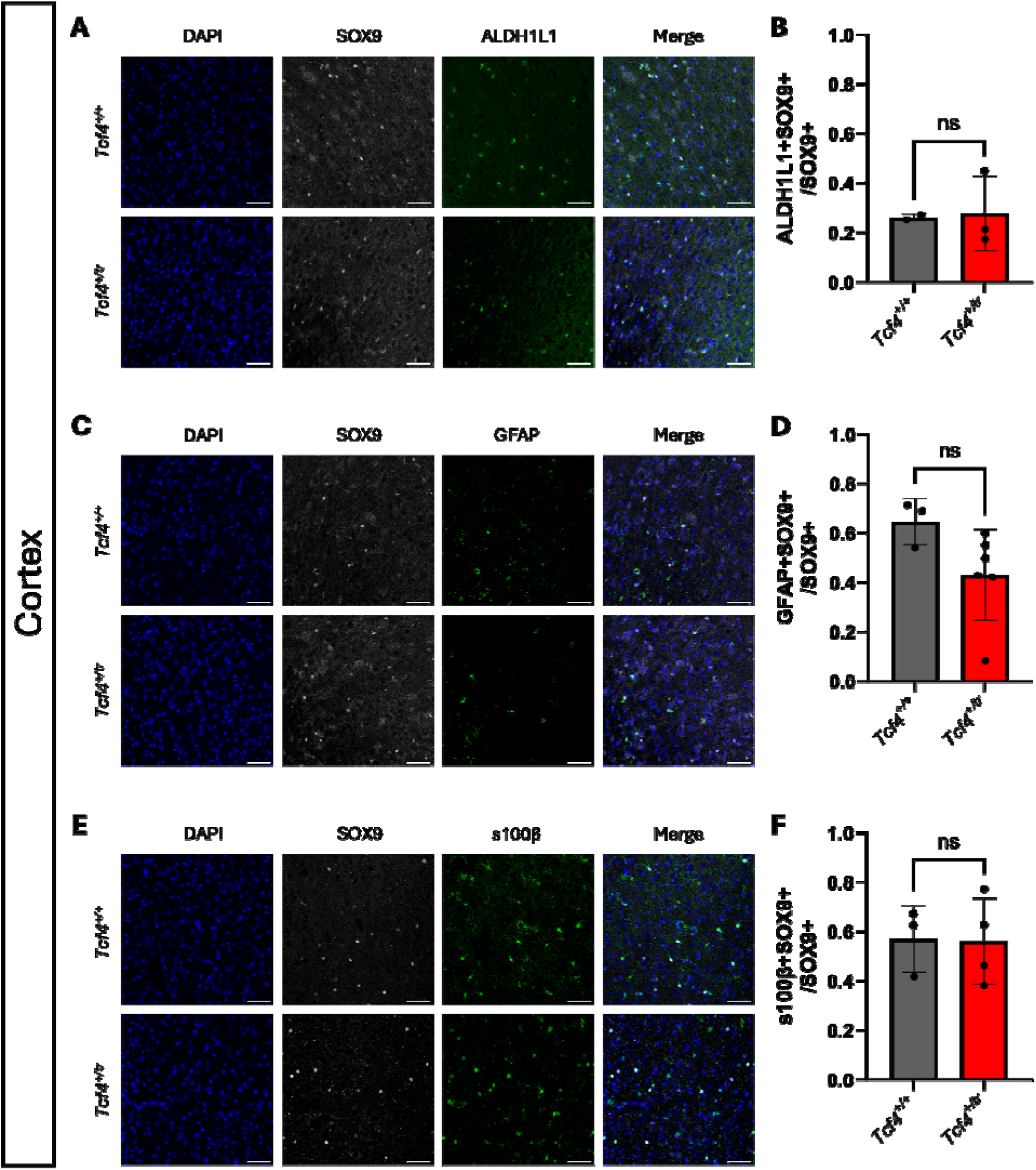
Astrocyte subpopulations in the cortex are unimpaired in the PTHS mouse model. (**A**) IHC of the ALDH1L1+ astrocytes in the P42 cortex of *Tcf4^+/tr^* and WT littermates. Tissue stained for DAPI (blue, column 1), SOX9 (gray, column 2), and ALDH1L1 (green, column 3). (**B**) Summary plot showing no change in the proportion of ALDH1L1+SOX9+/SOX9+ cells between *Tcf4^+/tr^* and WT mice (*Tcf4^+/tr^*0.2784 ± 0.08726 *n* = 3 mice, WT 0.2620 ± 0.009973 *n* = 2 mice; *P* = 0.8932). (**C**) IHC of the fibrous astrocyte population in the P42 cortex of *Tcf4^+/tr^* and WT littermates. Tissue stained for DAPI (blue, column 1), SOX9 (gray, column 2), and GFAP (green, column 3). (**D**) Summary plot showing no change in the proportion of GFAP+SOX9+/SOX9+ cells between *Tcf4^+/tr^* and WT mice (*Tcf4^+/tr^* 0.4302 ± 0.07493 *n* = 6 mice, WT 0.6474 ± 0.05448 *n* = 3 mice; *P* = 0.1016). (**E**) IHC of the protoplasmic astrocyte population in the P42 cortex of *Tcf4^+/tr^* and WT littermates. Tissue stained for DAPI (blue, column 1), SOX9 (gray, column 2), and s100β (green, column 3). (**F**) Summary plot showing no change in the proportion of s100β+SOX9+/SOX9+ cells between *Tcf4^+/tr^* and WT mice (*Tcf4^+/tr^* 0.5610 ± 0.08728 *n* = 4 mice, WT 0.5718 ± 0.07874 *n* = 3 mice, *P* = 0.9329). Data are presented as mean values ± SEM, scale bars 50 µm.

**Figure 4:**
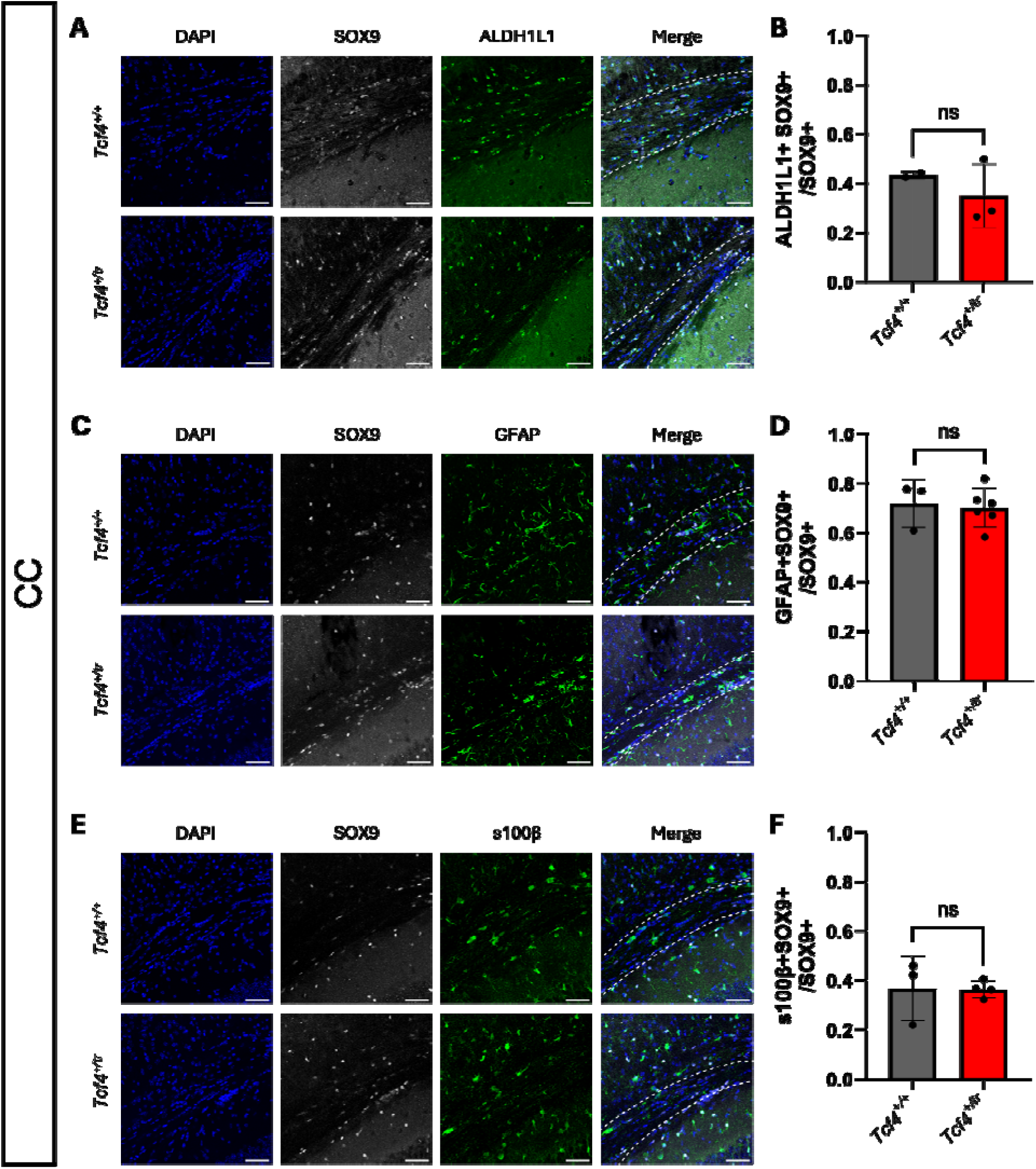
Astrocyte subpopulations in the CC are unimpaired in the PTHS mouse model. (**A**) IHC of ALDH1L1+ astrocytes in the P42 CC of *Tcf4^+/tr^* and WT littermates. Tissue stained for DAPI (blue, column 1), SOX9 (gray, column 2), and ALDH1L1 (green, column 3). (**B**) Summary plot showing no change in the proportion of ALDH1L1+SOX9+/SOX9+ cells between *Tcf4^+/tr^* and WT mice (*Tcf4^+/tr^* 0.3503 ± 0.07464 *n* = 3 mice, WT 0.4342 ± 0.009748 *n* = 2 mice; *P* = 0.4489). (**C**) IHC of the fibrous astrocyte population in the P42 CC of *Tcf4^+/tr^* and WT littermates. Tissue stained for DAPI (blue, column 1), SOX9 (gray, column 2), and GFAP (green, column 3). (**D**) Summary plot showing no change in the proportion of GFAP+SOX9+/SOX9+ cells between *Tcf4^+/tr^* and WT mice (*Tcf4^+/tr^* 0.7018 ± 0.03208 *n* = 6 mice, WT 0.7185 ± 0.05551 *n* = 3 mice; *P* = 0.7875). (**E**) IHC of the protoplasmic astrocyte population in the P42 CC of *Tcf4^+/tr^* and WT littermates. Tissue stained for DAPI (blue, column 1), SOX9 (gray, column 2), and s100β (green, column 3). (**F**) Summary plot showing no change in the proportion of s100β+SOX9+/SOX9+ cells between *Tcf4^+/tr^* and WT mice (*Tcf4^+/tr^* 0.3638 ± 0.01704 *n* = 4 mice, WT 0.3671 ± 0.07487 *n* = 3 mice; *P* = 0.9621). Data are presented as mean values ± SEM, scale bars 50 µm.

It was previously shown that conditional deletion of Tcf4 only in the ventral proliferative zones using the Nkx2.1-cre::*Tcf4*^fl/fl^ mouse resulted in a population of ventrally-derived “ectopic astrocytes” that had errantly migrated into the dorsal cortex by P15 and were continuously identifiable by P60 ^16^. We were interested in determining if misallocation of ventrally-derived astrocytes was associated with germline heterozygous mutations in *Tcf4*, which more closely models PTHS ^8^. We crossed Nkx2.1-Cre with a conditional TdTomato reporter line (TdTom) and generated Nkx2.1::TdTom reporter mice. These reporter mice were then crossed with *Tcf4^+/tr^*mice to quantify ventrally-derived astrocyte populations in WT and *Tcf4^+/tr^*mice. We performed IHC for SOX9 on P28 Nkx2.1::TdTom::*Tcf4^+/tr^*and WT littermates and quantified the overall population of ventrally-derived astrocytes in the cortex and CC. Consistent with our results comparing *Tcf4^+/tr^* and WT littermates (Figure 2), we observed no overall difference in the proportion of SOX9+DAPI+/DAPI+ cells regardless of their origin between genotypes in either the cortex (Supp Figure 1A, B) or CC (Supp Figure 1C, D). Similarly, looking at subtypes of astrocytes, we observed no difference in the proportions of GFAP+SOX9+ or s100β+SOX9+ cells in either the cortex (Figure 5) or CC (Figure 6). To determine if germline heterozygous mutation of *Tcf4* resulted in mislocalization of ventrally-derived astrocytes, we searched for SOX9+TdTom+ cells in the dorsal cortex, but found no such cells. Within sections containing the CC, we identified a single SOX9+TdTom+ cell that was located below the CC in a single *Tcf4^+/tr^* animal, however, it was difficult to differentiate which of two neighboring DAPI+ cells was TdTom+, making it difficult to identify the cell as an astrocyte (Supp Figure 2).

**Figure 5:**
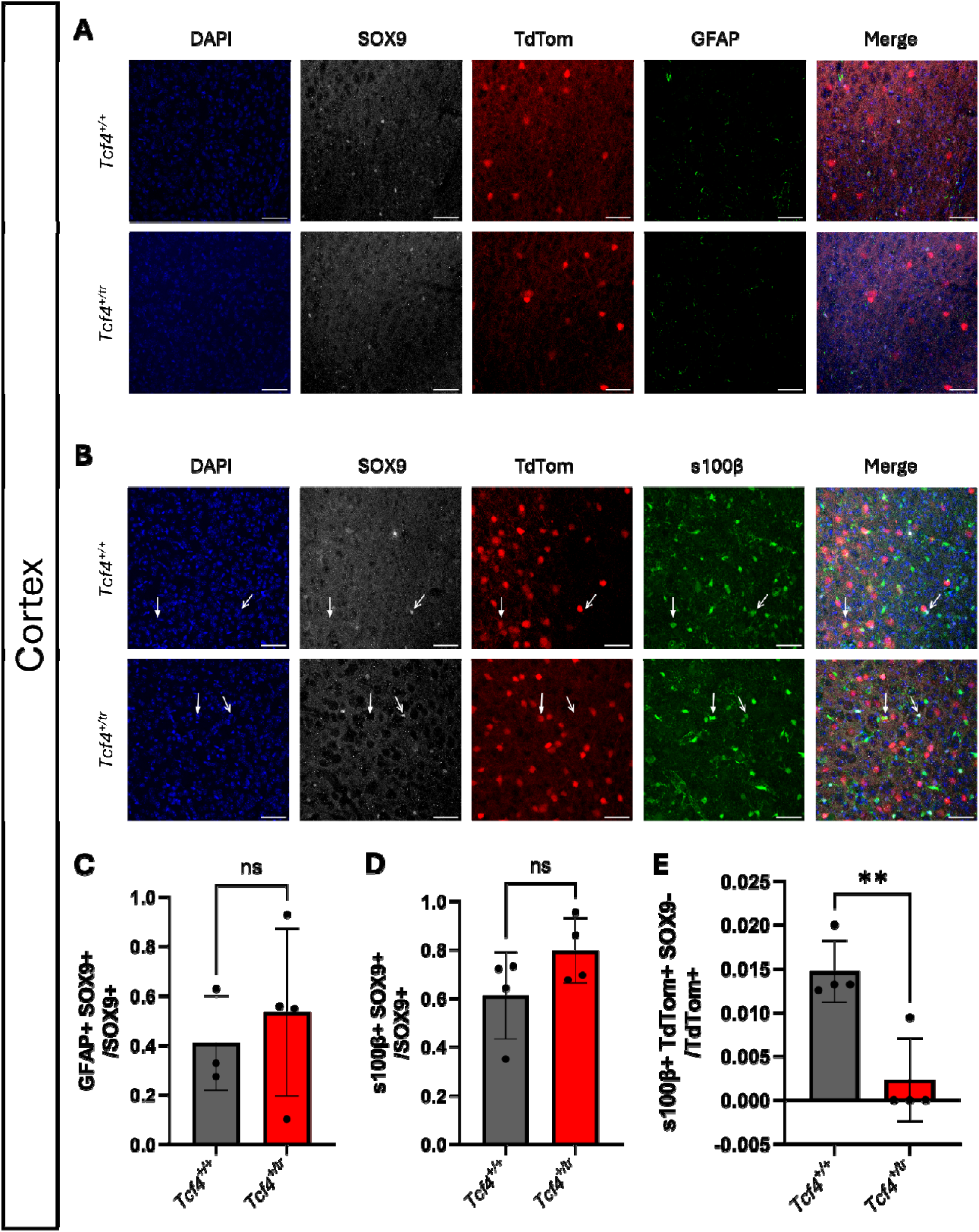
Quantification of ventrally-derived protoplasmic astrocyte populations in the PTHS mouse model. (**A**) IHC of the ventrally-derived GFAP+ astrocyte population in the dorsal cortex of P28 Nkx2.1::TdTom::*Tcf4^+/tr^* and WT littermates. Tissue stained for DAPI (blue, column 1), SOX9 (gray, column 2), TdTom (red, column 3), and GFAP (green, column 4). (**B**) IHC of the ventrally-derived protoplasmic s100β+ astrocyte population in the cortex of P28 Nkx2.1::TdTom::*Tcf4^+/tr^* and WT littermates. Tissue stained for DAPI (blue, column 1), SOX9 (gray, column 2), TdTom (red, column 3), and s100β (green, column 4). Closed arrows identify s100β+TdTom+SOX9-cells while open arrows identify s100β+TDT-SOX9+ cells. (**C**) Summary plot showing no change in the proportion of GFAP+SOX9+/SOX9+ cells in the cortex between *Tcf4^+/tr^* and WT mice (*Tcf4^+/tr^* 0.5337 ± 0.1695 *n* = 4 mice, WT 0.4095 ± 0.1106 *n* = 3 mice; *P* = 0.5984). No GFAP+TdTom+ cells were found in the cortex from either genotype. (**D**) Summary plot showing no change in the proportion of s100β+SOX9+/SOX9+ cells in the cortex between *Tcf4^+/tr^* and WT mice (*Tcf4^+/tr^* 0.7963 ± 0.06729 *n* = 4 mice, WT 0.6119 ± 0.08964 *n* = 4 mice; *P* = 0.1511). (**E**) Summary plot showing a significant difference in the proportion of s100β+TdTom+SOX9-/TdTom+ cells in the cortex between *Tcf4^+/tr^* and WT mice (*Tcf4^+/tr^*0.002359 ± 0.002359 *n* = 4 mice, WT 0.01470 ± 0.001772 *n* = 4 mice, *P* = 0.0058). Data are presented as mean values ± SEM, scale bars 50 µm.

**Figure 6:**
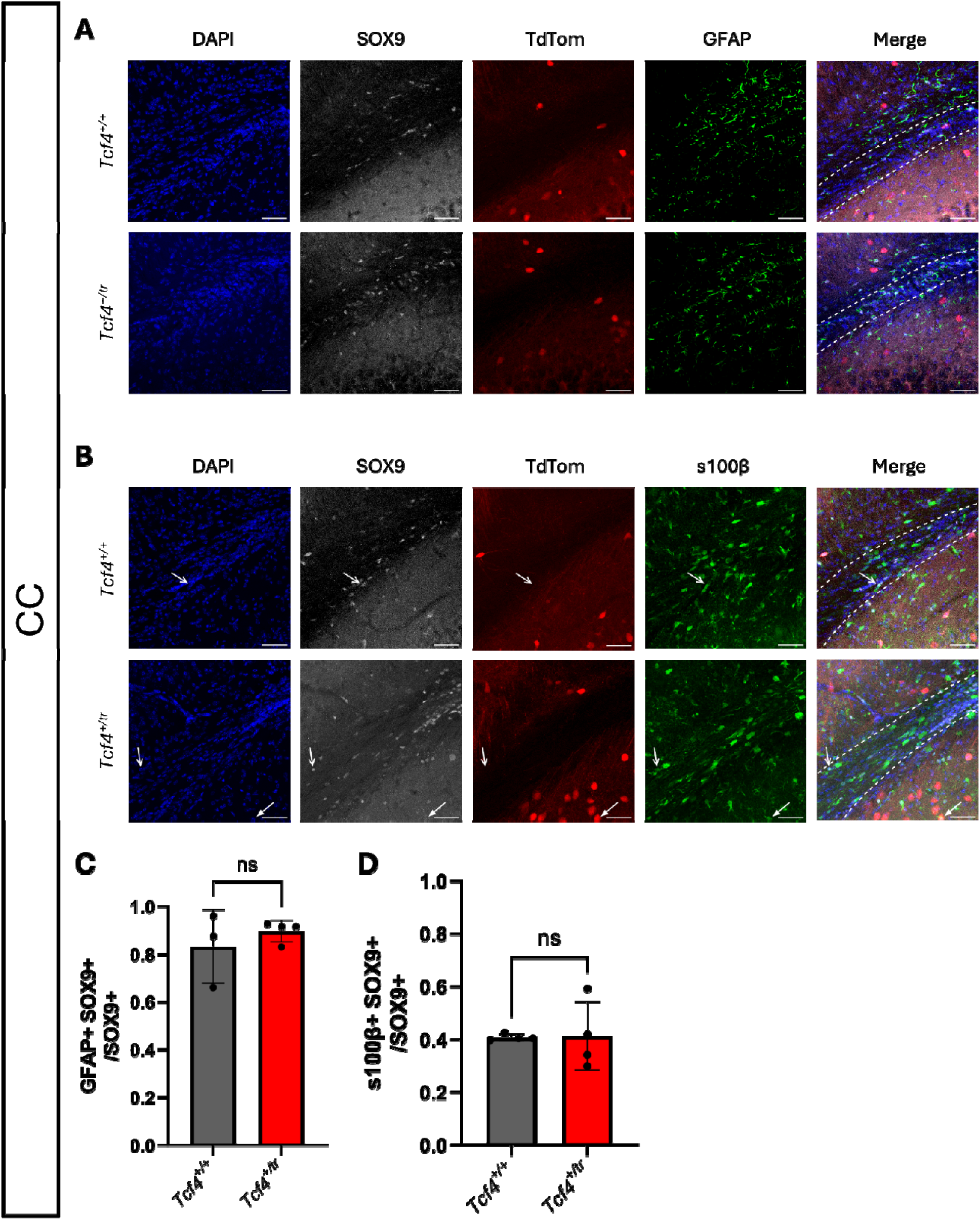
Characterization of ventrally-derived fibrous astrocyte populations in the PTHS mouse model. (**A**) IHC of the ventrally-derived GFAP+ astrocyte population in the CC of P28 Nkx2.1::TdTom::*Tcf4^+/tr^* and WT littermates. Tissue stained for DAPI (blue, column 1), SOX9 (gray, column 2), TdTom (red, column 3), and GFAP (green, column 4). (**B**) IHC of the ventrally-derived s100β+ protoplasmic astrocyte population in the CC of P28 Nkx2.1::TdTom::*Tcf4^+/tr^*and WT littermates. Tissue stained for DAPI (blue, column 1), SOX9 (gray, column 2), TdTom (red, column 3), and s100β (green, column 4). Closed arrows identify s100β+TdTom+SOX9-cells while open arrows identify s100β+TDT-SOX9+ cells. (**C**) Summary plot showing no change in the proportion of GFAP+SOX9+/SOX9+ cells in the CC between *Tcf4^+/tr^* and WT mice (*Tcf4^+/tr^* 0.8965 ± 0.02253 *n* = 4 mice, WT 0.8311 ± 0.08911 *n* = 3 mice; *P* = 0.4462). No GFAP+TdTom+ cells were found in the CC from either genotype. (**D**) Summary plot showing no change in the proportion of s100β+/SOX9+ cells in the CC between *Tcf4^+/tr^* and WT mice (*Tcf4^+/tr^*0.4124 ± 0.06480 *n* = 4 mice, WT 0.4068 ± 0.006267 *n* = 4 mice; *P* = 0.9342, 0.005603 ± 0.06510]. No s100β+TdTom+ cells were found in the CC from either genotype. Data are presented as mean values ± SEM, scale bars 50 µm.

Next, we searched for the presence of ventrally-derived fibrous and protoplasmic astrocyte subpopulations in the dorsal cortex of the Nkx2.1::TdTom::*Tcf4^+/tr^* model. We did not observe any cells that were clearly GFAP+TdTom+ in the dorsal cortex (Figure 5) or CC (Figure 6) of either *Tcf4^+/tr^*or WT mice. We did observe some TdTom+ cells that were contacted by GFAP processes, but these processes did not appear to be originating from the TdTom+ cells, and were instead likely processes coming off nearby astrocytes. We next searched for ventrally-derived protoplasmic astrocytes (s100β+TdTom+) in the dorsal cortex and in this case we were able to identify a few extremely rare examples of s100β+TdTom+ cells in the dorsal cortex. However, these cells were clearly negative for SOX9, suggesting they were myelinating OLs ^26^. Overall, we found a total of four s100β+TdTom+SOX9-cells in the dorsal cortex of WT mice, and only one such cell in the dorsal cortex of *Tcf4^+/tr^* mice. Although these counts were found to be significantly different by genotype (Figure 5E, Supp Figure 3), the overall rarity of this cell population greatly diminishes their potential relevance to pathology. No s100β+TdTom+ cells were found within the CC, though we did identify a single s100β+TdTom+ cell that was located below the CC in a single *Tcf4^+/tr^* animal (Figure 6B).

Overall, our findings indicate that heterozygous LOF mutations in *Tcf4* do not appear to disrupt the proportions of fibrous and protoplasmic astrocytes in the cortex and CC. In addition, germline *Tcf4* mutation did not appear to disrupt the allocation and localization of ventrally-derived astrocytes, as they remain in the ventral cortex. Therefore, at the cell population level, it does not appear that *Tcf4* mutations that lead to PTHS are altering the astrocyte lineage.

## DISCUSSION

Astrocytes are implicated in a variety of neurodegenerative and neuropsychiatric disorders including multiple sclerosis, Alzheimer disease, Parkinson disease, Huntington disease, and neuropsychiatric disorders ^27^. Emerging evidence suggests that ASD risk genes functionally converge on the regulation of neural progenitor cell proliferation and differentiation during corticogenesis, and the diverse functional roles of astrocytes contributing to regulating brain development and synaptic function implicates their association with ASD etiology. However, evidence for a role of astrocytes in ASD pathophysiology is relatively scarce, with a few postmortem brain studies identifying alteration in the number of astrocytes in ASD brain compared to controls ^18^, and a single nuclear RNA sequencing study identified evidence for an elevated state of activation in glial cell types including astrocytes ^28^.

Genome-wide association studies and whole exome sequencing studies continually identify genetic variants that associate the *TCF4* locus with a variety of psychiatric disorders including schizophrenia, bipolar, post-traumatic stress disorder, major depressive disorder, and ASD^29–34^. The most direct clinical association for TCF4 is PTHS, which is caused by autosomal dominant mutations in TCF4, and is characterized as a profound neurodevelopmental disorder ^8^. Mouse and human models of PTHS have consistently identified dysregulation in neural progenitor cell proliferation and specification in both dorsal and ventral proliferative zones during cortical development which leads to abnormal production of neuronal and glial lineages ^9,9–15^. Two studies are related to dysfunction in the astrocyte lineage and both only show abnormalities due to homozygous deletion of *Tcf4* ^10^, which in mice results in death around birth ^35^. Mesman and colleagues showed Tcf4^-/-^ mice at embryonic stages were devoid of midline glia (GFAP+) in the indusium griseum and callosal wedge, which likely contributed to abnormal axon guidance during CC development, however this population of GFAP+ glia were not disrupted in Tcf4^+/-^ mice ^10^. In addition, when Tcf4 was conditionally deleted in the Nkx2.1 lineage (Nkx2.1-cre::Tcf4^fl/fl^), ventrally-derived protoplasmic and fibrous astrocytes were both ectopically observed in the dorsal cortex, however the overall numbers of astrocytes was unchanged ^16^. Together, these studies may suggest a role for TCF4 in astrocyte migration, which matches prior studies showing TCF4 regulation of neural progenitor and neuronal migration ^9,36–38^.

In our survey of astrocytes in the PTHS mouse model, we did not observe any effect of TCF4 LOF on the proportions of fibrous or protoplasmic astrocytes, even though we observed expression of TCF4 in the astrocyte lineage. Moreover, in this context, we did not observe ectopic allocation of ventrally-derived astrocytes in the dorsal cortex. This result is somewhat surprising given that both the neuronal and oligodendroglial lineage are affected by heterozygous TCF4 LOF, but may be due to the homeostatic nature of astrocytes and their ability to proliferate, which may contribute to maintaining appropriate astrocyte density. This compensatory mechanism appears to be at work in the Nkx2.1-cre::Tcf4^fl/fl^ model, as there is no difference in the overall density of astrocytes in the dorsal cortex even though ventrally-derived astrocytes are misallocated to the dorsal cortex ^16^. We have previously observed that TCF4 LOF disrupts the proportions of specific subsets of inhibitory neurons and oligodendrocytes, and in these same cell populations we observed enrichment of differentially expressed genes (DEGs), however this enrichment of DEGs was not observed in astrocytes ^12,13^. Our results do not completely rule out a role for TCF4 in regulating astrocyte function, as TCF4 is clearly expressed in astrocytes and we only surveyed astrocytes at the cell population level. Future studies will need to dig deeper into astrocyte gene expression and function to determine if dysregulation of TCF4 is consequential for astrocyte biology.

In summary, our results indicate that in the context of PTHS, TCF4 does not disrupt the proportion of fibrous and protoplasmic astrocytes in the murine dorsal cortex. Moreover, it does not appear that germline heterozygous TCF4 LOF leads to misallocation of ventrally-derived astrocytes into the dorsal cortex. Our results predict that at the cell population level, the density of astrocytes are not disrupted in PTHS.

## FIGURE LEGENDS

**Supplemental Figure 1:**
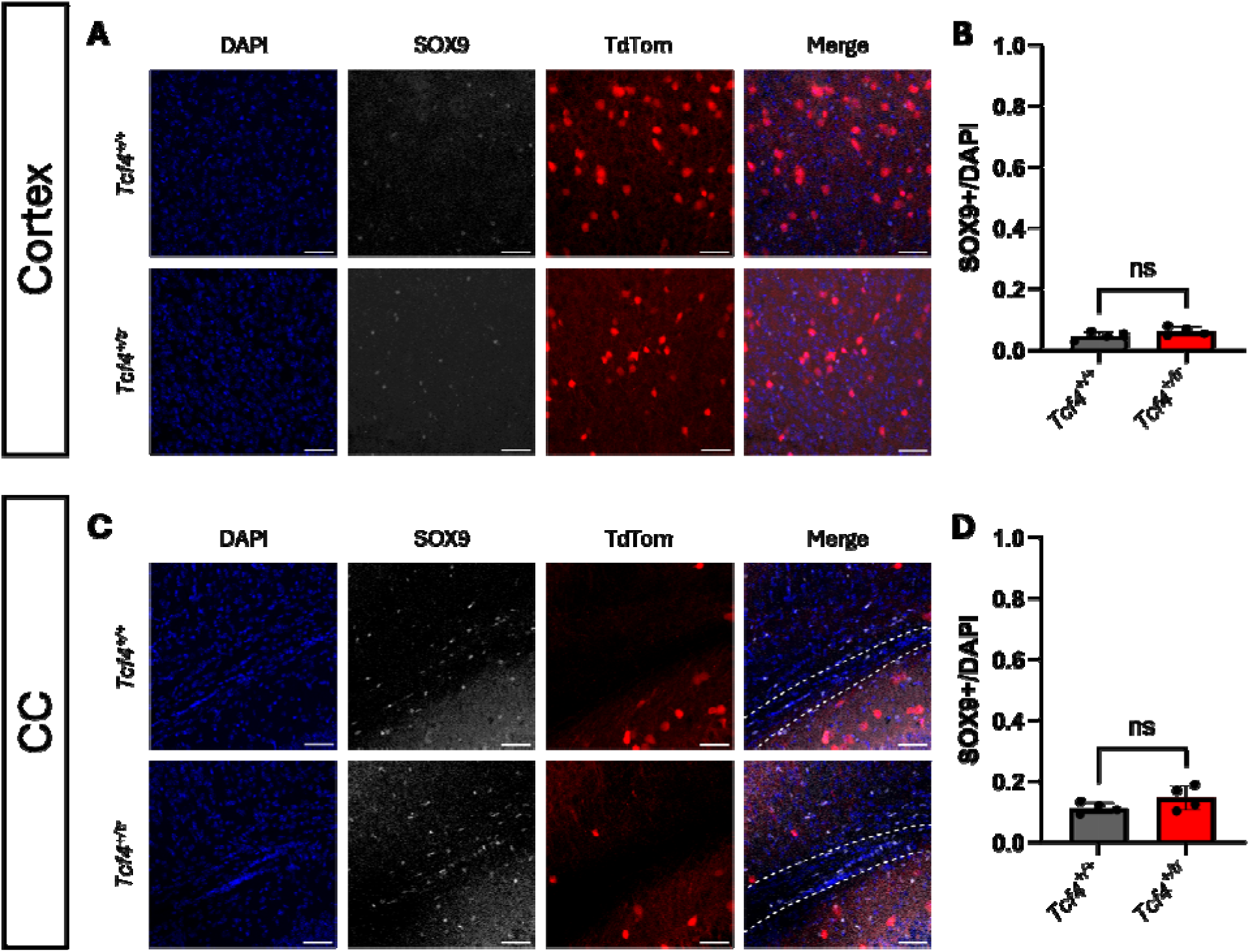
Quantification of ventrally-derived astrocyte populations in the cortex and CC of the PTHS mouse model. (**A**) IHC of the total ventrally-derived astrocyte population in the cortex of P28 Nkx2.1::TdTom::*Tcf4^+/tr^* and WT littermates. Tissue stained for DAPI (blue, column 1), SOX9 (gray, column 2), and TdTom (red, column 3). (**B**) Summary plot showing no change in the proportion of SOX9+DAPI+/DAPI+ cells between *Tcf4^+/tr^* and WT mice (*Tcf4^+/tr^* 0.06183 ± 0.007300 *n* = 4 mice, WT 0.04662 ± 0.006277 *n* = 4 mice, *P* = 0.1654). No cortical SOX9+TdTom+ cells were found in either genotype. (**C**) IHC of the total ventrally-derived astrocyte population derived in the CC of P28 Nkx2.1::TdTom::*Tcf4^+/tr^*and WT mice. Tissue stained for DAPI (blue, column 1), SOX9 (gray, column 2), and TdTom (red, column 3). (**D**) Summary plot showing no change in the proportion of SOX9+DAPI+/DAPI+ cells between *Tcf4^+/tr^* and WT mice (*Tcf4^+/tr^* 0.1455 ± 0.02022 *n* = 4 mice, WT 0.1124 ± 0.009220 *n* = 4 mice, *P* = 0.1866). No SOX9+TdTom+ cells were found in the CC or surrounding tissue in either genotype. Data are presented as mean values ± SEM, scale bars 50 µm.

**Supplemental Figure 2:**
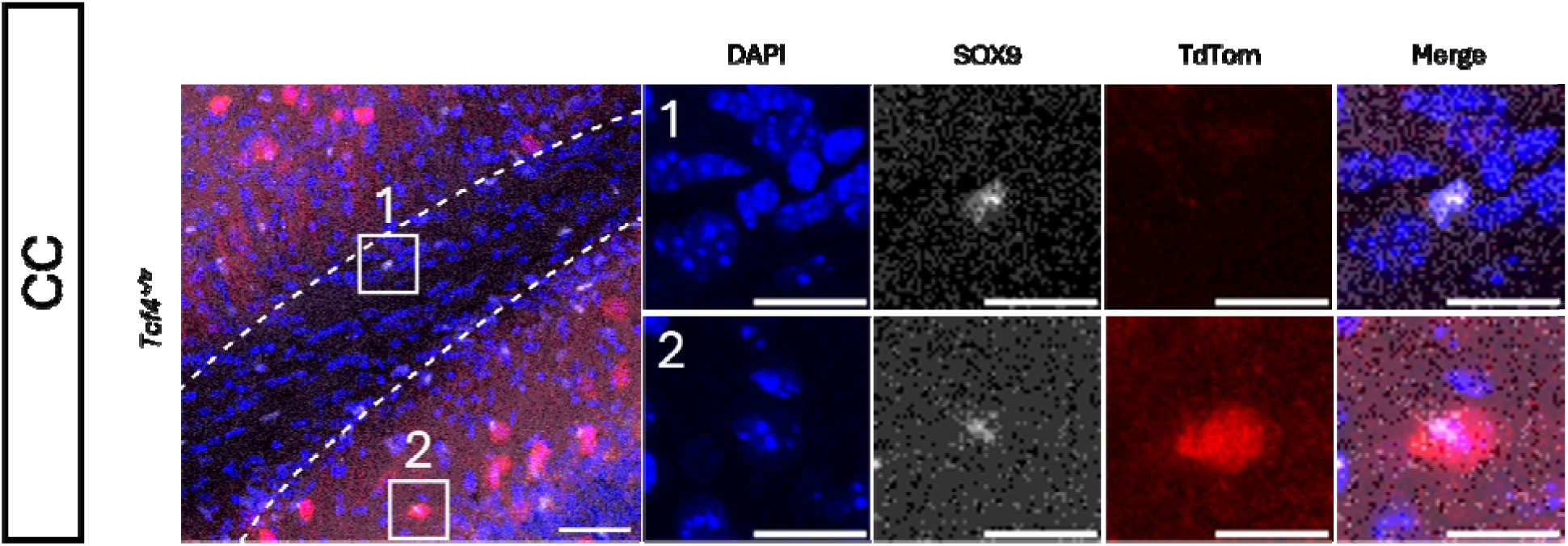
Magnified view of confocal imaging for SOX9 and TdTom. Merged IHC image of the CC of a *Tcf4^+/tr^* mouse. Scale bar 50 µm (Left panel). The cell numbered 1 depicts a SOX9+TdTom-astroglia. The cell numbered 2 depicts a possible SOX9+TdTom+ cell. Scale bars 10 µm.

**Supplemental Figure 3:**
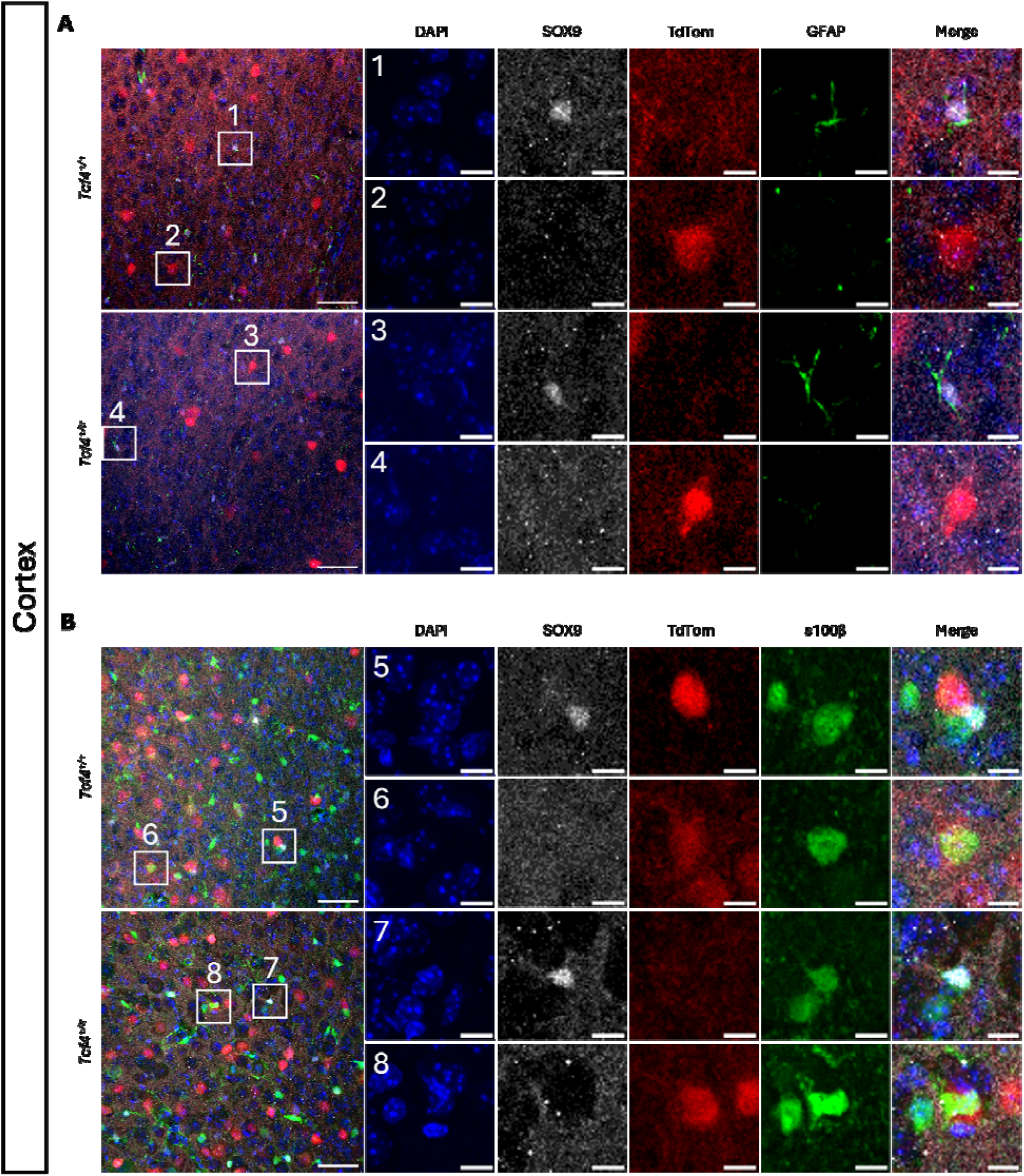
Magnified view of confocal imaging for astrocyte markers and TdTom reporter. Merged IHC images of the cortex from Figures 5 and 6. Scale bars 50 µm. (**A**) Cells numbered 1 and 3 depict SOX9+TdTom-GFAP+ astroglia. Cells numbered 2 and 4 depict SOX9-TdTom+GFAP-cells. (**B**) Cells numbered 5 and 7 depict SOX9+TdTom-s100β+ astroglia. Cells numbered 6 and 8 depict SOX9-TdTom+s100β+ cells, likely to be myelinating OLs. Scale bars 10 µm.

## AUTHOR CONTRIBUTION

**S. Stump:** Methodology, Investigation, Validation, Formal analysis, Data curation, Visualization, Writing -Original Draft, Writing - Review & Editing. **J.F. Bohlen:** Methodology, Investigation, Data curation, Supervision, Writing - Original Draft, Writing - Review & Editing. **B. Phan:** Methodology, Investigation, Data curation, Formal Analysis, Visualization. **B.J. Maher:** Conceptualization, Methodology, Resources, Data curation, Visualization, Writing - Original Draft, Writing - Review & Editing, Supervision, Project administration, Funding acquisition.

## ACKNOWLEDGEMENTS

We are grateful for the vision and generosity of the Lieber and Maltz families, who made this work possible. This project was supported by the Lieber Institute for Brain Development, the National Institute of Mental Health (Grant No. R01MH110487 to BJM), and the Pitt-Hopkins Research Foundation and Penn Medicine Orphan Disease Center (Grant No. MDBR-24-002-PittHopkins to BJM). The content is solely the responsibility of the authors and does not necessarily represent the official views of the National Institutes of Health. Figure panel 1B and 2A created in BioRender. Maher, B. (2025) https://BioRender.com/nh8cyou

